# Long-term root electrotropism reveals habituation and hysteresis

**DOI:** 10.1101/2023.07.28.551054

**Authors:** Maddalena Salvalaio, Giovanni Sena

## Abstract

Plant roots sense many physical and chemical cues in soil, such as gravity, humidity, light and chemical gradients, and respond by redirecting their growth towards or away from the source of the stimulus. This process is called tropism. While gravitropism is the tendency to follow the gravitational field downwards, electrotropism is the alignment of growth with external electric fields and the induced ionic currents. Although root tropisms are at the core of their ability to explore large volumes of soil in search of water and nutrients, the molecular and physical mechanisms underlying most of them remain poorly understood. We have previously provided a quantitative characterization of root electrotropism in Arabidopsis (*Arabidopsis thaliana*) primary roots exposed for 5 hours to weak electric fields, showing that auxin asymmetric distribution is not necessary but that cytokinin biosynthesis is. Here, we extend that study showing that long-term electrotropism is characterized by a complex behavior. We describe overshoot and habituation as key traits of long-term root electrotropism in Arabidopsis and provide quantitative data about the role of past exposures in the response to electric fields (hysteresis). On the molecular side, we show that cytokinin, although necessary for root electrotropism, is not asymmetrically distributed during the bending.

Overall, the data presented here represent a significant step forward towards the understanding of the molecular mechanisms regulating electrotropism in plants and provide a quantitative platform for future studies on the genetics of this and other tropisms.

## INTRODUCTION

Plant roots exhibit remarkable adaptive behavior by navigating the complex soil environment in search of water and nutrients. This relies on the ability to modify the direction of growth in response to external stimuli, such as water potential, gravity, light, temperature, mechanical pressure, electric charges, and chemical concentrations, a phenomenon generally known as tropism (Izzo and Aronne, 2021). Although significant progress has been made in understanding the molecular aspects of a few root tropisms (Muthert et al., 2020), there are still several key sensing mechanisms that remain elusive.

Physical and chemical modifications of soil, derived from its geological formation, give rise to mineral ions essential for plant metabolism (Pozdnyakov and Pozdnyakova, 2002), while organic ions are released by living organisms in soil (England and Robert, 2022) (England and Robert, 2022). The non-uniform distribution of these ions in soil contributes to the spontaneous formation of local, weak, electric fields (Pozdnyakov and Pozdnyakova, 2002). Additionally, microorganisms and plant roots generate their own bioelectric fields that contribute to the complex and dynamic electromagnetic signature of soil (Takamura, 2006; Chabert et al., 2015). These transient electric fields and their associated currents in soil potentially encode valuable information regarding the localization of water, micronutrients, and organisms, offering a selective advantage to those capable of sensing them.

Root electrotropism, also known as galvanotropism, is the ability of plant roots to sense and grow towards or away electric charges, or to align with local electric fields and ionic currents (Navez, 1927). First documented in 1882 by Elfving (Elfving, 1882) and later rediscovered in the early 20th century by Ewart and Bayliss (Ewart and Bayliss, 1905), root electrotropism has been sporadically studied in maize (*Zea mays*), peas (*Pisum sativum*), and beans (*Vigna mungo*), yielding conflicting results (Wolverton et al., 2000). Recently, we have provided a quantitative characterization of the short-term electrotropic response of *Arabidopsis* primary roots in the first 5 hours of exposure (Salvalaio et al., 2022). Bending of individual root tips in electric fields occurs in minutes to a few hours (Salvalaio et al., 2022), but little is known about long-term behavior.

Interestingly, some sensing processes in animals exhibit peculiar kinetics labeled as “non-associative learning”, such as habituation and memory, emerging only after long or repeated exposure to the stimulus (Thompson and Spencer, 1966; Rankin et al., 2009). Although some attempts have been made to study habituation and stress priming (a form of memory) in non-neuronal organisms such as slime molds (Boisseau et al., 2016) and plants (Bruce et al., 2007; Crisp et al., 2016; Hilker and Schmülling, 2019) and to generalize its theoretical foundations (Bonzanni et al., 2019) it remains unclear to which extent these are phenomena requiring the complexity of animal nervous systems or, instead, are fundamental and unavoidable traits of any sensing mechanism.

In this work, we extend our previous analysis of root electrotropism in Arabidopsis (Salvalaio et al., 2022) and reveal a long-term behavior reminiscent of habituation and hysteresis when roots are exposed to a comparable electric field for over 17 hours. This longer duration of the stimulus would model natural scenarios where the physical or biological source of the electric field in soil is persistent.

By characterizing the anatomical, molecular, and kinetic aspects of this phenomenon, we strive to bridge the gap between plant and animal electric sensing mechanisms, contributing to our understanding of the evolutionary adaptations employed by organisms in response to their environment.

## RESULTS

### Long-term root electrotropism results in an overshoot followed by a new set-point angle

To image the long-term behavior of Arabidopsis roots exposed to an external electric field, we adapted the root electrotropism assay previously described (Salvalaio et al., 2022) to conduct experiments in sterile conditions (details in Materials and Methods). Briefly, the setup includes a small and transparent box wired as an electrolytic cell (V-box), where 7-day-old seedlings are mounted vertically in a buffered liquid medium and between two foil electrodes connected to a power supply to maintain a constant difference of electric potential (Supplemental Figure S1, a). This results in a “nominal” electric field EF and an estimated current Ir impacting the root (Table 1 and Material and Methods for details). Throughout the experiments, 3.6 l of the liquid medium is circulated from a reservoir through the V-box to minimize any electrochemical gradient and to maintain a constant temperature. The entire circulation system is closed and sterile. A Raspberry Pi camera is positioned to take images of the growing roots every 0.5 to 1.0 h, and the angle between each root tip and the gravity vector is measured (Supplemental Figure S1, b).

**Table 1.** Nominal electric field and corresponding current impacting the root. Nominal electric field (EF) and mean ± standard deviation of the measured *I*_*r*_ = *I* × *A*_*r*_/*A*_*e*_, where *I* is the total current intensity measured in the circuit, *A*_*r*_ is the estimated area of the root surface facing each electrode, *A*_*e*_ is the surface area of each electrode.

| **SET-UP** | **EF (V/cm)** | **$I_r$ (mA)** |
| --- | --- | --- |
| V-box | 1.0 | $0.16 \pm 0.03$ |
| V-box | 1.5 | $0.25 \pm 0.04$ |
| V-box | 2.0 | $0.40 \pm 0.08$ |
| V-box | 2.5 | $0.45 \pm 0.09$ |
| V-box | 3.0 | $0.60 \pm 0.12$ |
| V-slide | 3.0 | $0.48 \pm 0.10$ |

As we have previously shown (Salvalaio et al., 2022), the early response of Arabidopsis primary roots exposed to a perpendicular EF of 1.5 V/cm is a reorientation of their growth vector (tropism) toward the negative electrode (cathode) (t-test between 1.5 and 0 V/cm at 3 h, p < 0.001) (Figure 1, a). However, this orientation is not maintained over time and after about 3 hours the root tips slowly bend downwards until they reach a new stable angle (Figure 1, a). Overall, this long-term response can then be described as an initial overshoot, followed by relaxation and steady-state, or set-point angle. Crucially, the set-point angle of roots exposed in the V-box for 17 h to 1.5 V/cm was significantly different from that of roots not exposed to the field (Figure 1, a; t-test between 1.5 and 0 V/cm at 17 h, p < 0.001).

**Figure 1.**
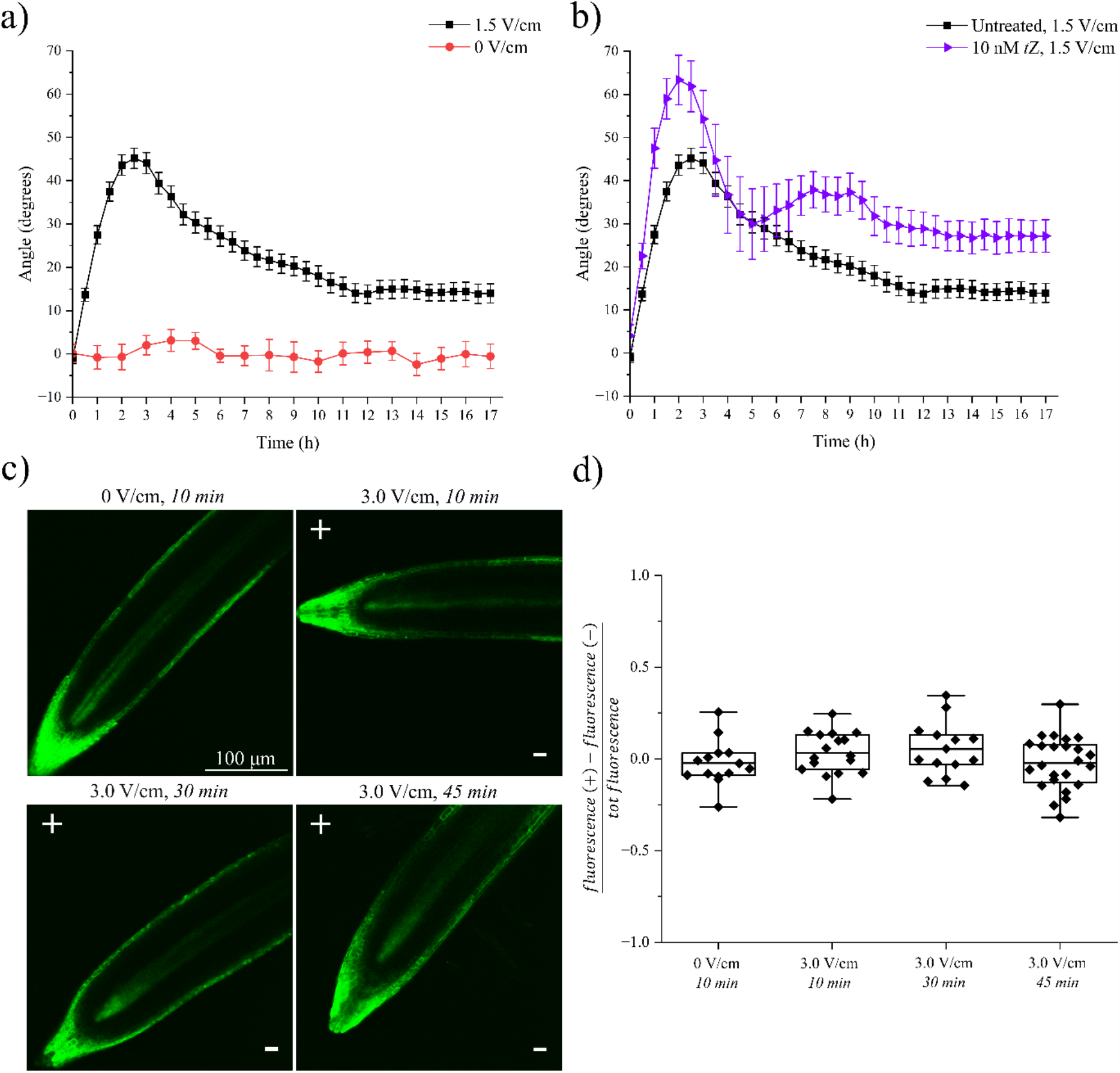
Long-term electrotropic response of Arabidopsis root and cytokinin distribution. **a**, Average root tip orientations relative to the gravity vector versus time at 0 and 1.5 V/cm; 0 V/cm, N=19, R=4; 1.5 V/cm, N=71, R=18. Error bars, s.e.m. **b**, Average root tip orientations relative to the gravity vector over time of plants exposed to 1.5 V/cm treated with 10 nM *t*Z (N=16, R=4) vs untreated (N=71, R=18). Error bars, s.e.m. **c**, Representative confocal images of Arabidopsis primary roots expressing the cytokinin-response reporter *TCSn::GFP* and exposed to EF for 10 to 45 minutes; the position of the positive and negative electrodes relative to the roots is indicated with + and -, respectively. Scale bar, 100 µm. **d**, Quantification of the symmetric distribution of GFP fluorescence in *TCSn::GFP* roots exposed to EF (the symmetry parameter is described in the main text); data points are individual roots; boxplots span the first to the third quartiles of the data, whiskers indicate minimum and maximum values and horizontal lines in the boxes represent the mean. See main text for statistical analysis. 0 V/cm, N=14, R=3; 3.0 V/cm for 10 min, N=17, R=6; 3.0 V/cm for 30 min, N=14, R=5; 3.0 V/cm for 45 min, N=24, R=7. N, sample size; R, number of replicates.

### Cytokinin is symmetrically distributed and is not depleted during root electrotropism

Previously, we have also shown that root electrotropism requires the biosynthesis of the cytokinin *trans*-zeatin (*t*Z) (Salvalaio et al., 2022). Specifically, we found that mutants of CYP735A1, a cytochrome P450 monooxygenase that catalyzes the biosynthesis of *t*Z (Takei et al., 2004), are not electrotropic but respond to the EF if supplied with 10 nM of *t*Z in the medium (Salvalaio et al., 2022). We wondered whether the mere depletion of available *tZ* in the tissue could explain the relaxation during electrotropism. To test this, we grew wild-type seedlings in the same medium but supplemented with 10 nM of *t*Z, reasoning that this would prevent a significant depletion in the tissue, and then exposed them to 1.5 V/cm for 17 h in the same medium. Nonetheless, relaxation was observed even in these conditions (Figure 1, b; t-test between the maximum at 2h and the plateau at 15-17 h, p < 0.001), indicating that *tZ* depletion is an unlikely explanation.

The hormone cytokinin, and specifically its asymmetric distribution, has also been associated with the variable root tip orientation in response to gravity, or Gravitropic Set-point Angle (GSA), measured in lateral roots (Waidmann et al., 2019), so we wondered whether the relaxation and new steady-state observed in response to EF are based on a similar mechanism. In other words, is the EF inducing a GSA through an asymmetric distribution of cytokinin? To test this hypothesis, we used a previously developed microscope chamber (Salvalaio et al., 2022) (V-slide, see Material and Methods) to expose roots expressing the cytokinin-sensitive reporter *TCSn::GFP* (Zürcher et al., 2013) to EF while imaging them on a confocal microscope. After exposing them to a 3.0 V/cm EF in the V-slide (Ir = 0.48 ± 0.10 mA, see Table 1), we imaged those roots actively bending towards the negative electrode and calculated a “symmetry parameter” defined as the difference between the average fluorescence at the side of the root facing the positive electrode and that at the side of the root facing the negative electrode, divided by their sum (total fluorescence), so that zero represents symmetric distribution. Interestingly, we did not detect any statistically significant asymmetric distribution of cytokinin (Figure 1, c and d), as confirmed by a one-way ANOVA to test for a significant difference between 0 V/cm and 3 V/cm for 10, 30 and 45 min, with p = 0.26, and a one sample t-test to test for a significant difference between the sample mean and the value zero for the symmetry parameter: 0 V/cm, p = 0.50; 3 V/cm for 10 minutes, p = 0.28; 3 V/cm for 30 minutes, p = 0.19; 3 V/cm for 45 minutes, p = 0.45. This new result, together with our previous data showing no role for auxin distribution and amyloplast-bearing root tips (Salvalaio et al., 2022), indicates that the long-term root reorientation observed in EF is not a modified gravitropic response, or a GSA, but an independent electrotropic response. For this reason, we called the new set-point angle an Electrotropic Set-point Angle (ESA) and defined it as the average of root tip orientation in the last 2 hours of exposure to EF in our long-term experiments.

### Response curve and anomalous cellular morphology in higher electric fields

The new steady-state ESA supposedly represents an equilibrium between gravitropism and electrotropism, so we asked whether an increased EF would result in a higher ESA.

We exposed roots to 1.0, 1.5, 2.0 and 2.5 V/cm in the V-box (see Table 1 for corresponding Ir) and found a remarkable linear correlation between EF and ESA, or response curve (Fig. 2, a; R^2^ = 0.99). We noticed that the variance in the ESA distribution at higher EF was getting larger, so we wanted to characterize better the response to strong EF. It has been previously suggested that roots can be damaged by high EFs and that this results in root tips pointing towards the anode (Stenz and Weisenseel, 1993). Indeed, we observed that, while 100% of roots bent towards the negative electrode when exposed to 1.0 V/cm, an increasing number of roots bent toward the positive electrode when exposed to higher electric fields and ionic currents (Figure 2, b). To investigate this further, we imaged with confocal microscopy in the V-slide roots exposed to 3.0 V/cm for 45 minutes and observed that, although cells in 64% (±7.4% s.e.p.) of the roots maintained their normal turgor, 19% (± 6.1% s.e.p.) and 17% (± 5.8% s.e.p.) of roots showed at least one cell in the cortical layer of the transition zone of the root facing the positive electrode significantly expanded or burst, respectively (Figure 2, c). Interestingly, all roots with burst cells did orient toward the positive electrode, confirming the previous hypothesis that this might be due to tissue damage and suggesting a simple mechanical recoil as the main mechanism.

**Figure 2.**
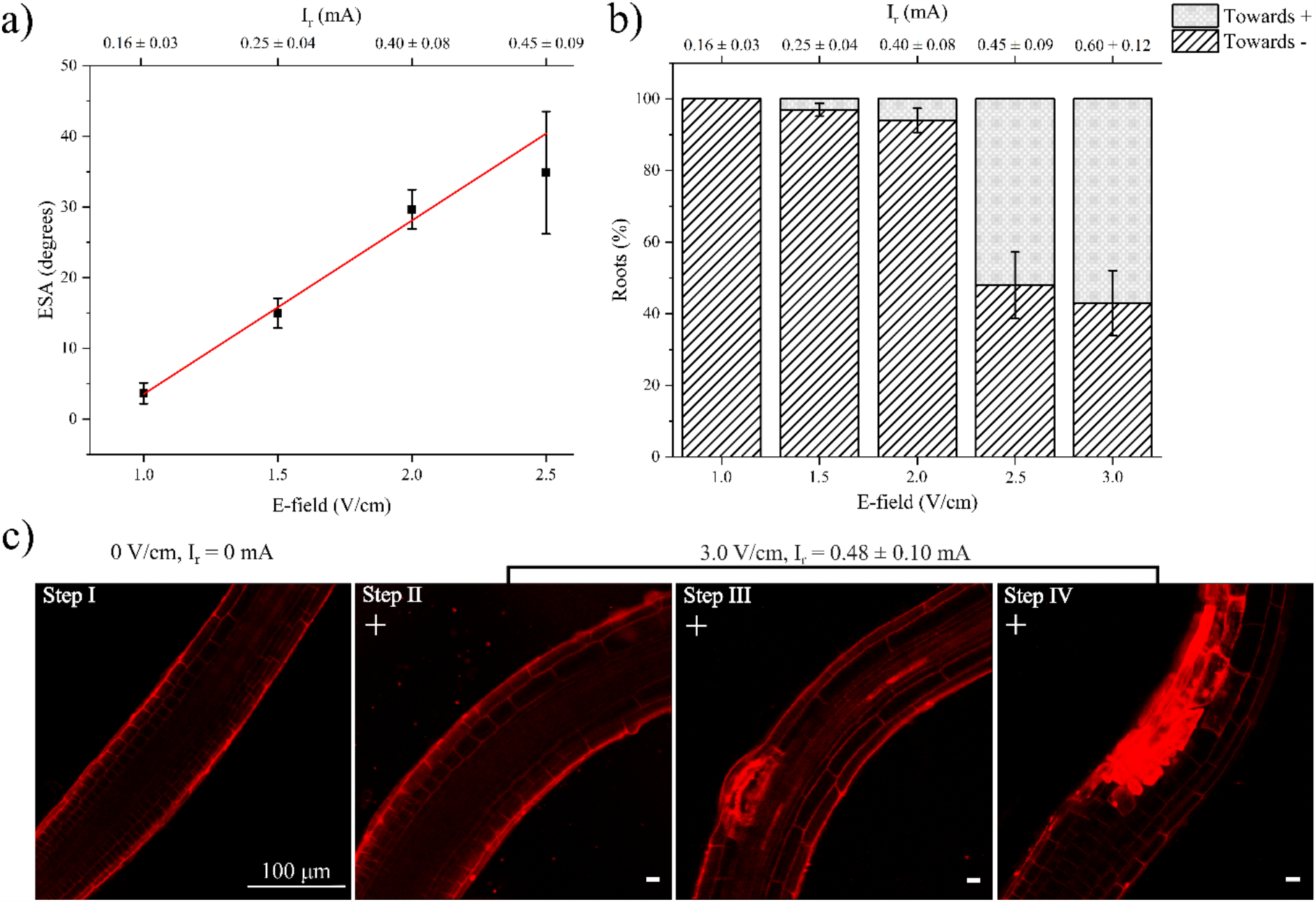
Long-term Arabidopsis root response to different E-field intensities. **a**, Long-term response curve: average ESA vs applied EF; 1.0 V/cm, N=34, R=7; 1.5 V/cm, N=75, R=18; 2.0 V/cm, N=47, R=11; 2.5 V/cm, N=14, R=5. Error bars, s.e.m. **b**, Percentage of roots orienting towards the positive and negative electrodes after exposure to different EFs; 1.0 V/cm, N=34, R=7; 1.5 V/cm, N=77, R=18; 2.0 V/cm, N=50, R=11; 2.5 V/cm, N=29, R=6; 3.0 V/cm, N=30, R=6. Error bars, s.e.p. **c**, Sequence of confocal images of representative roots exposed to 3.0 V/cm. Step I, before exposure; Step II, electrotropism towards the negative electrode; Steps III and IV, expansion and damage of cortical cell (arrow). Red channel, propidium iodide. Scale bar, 100 µm. 0 V/cm, N=14, R=3; 3.0 V/cm, N=42, R=12. N, sample size; R, number of replicates.

### The electrotropic set-point angle (ESA) is the result of habituation

The relaxation process described in the previous section suggests that the electrotropic response is weakening after about 3 h (duration of the overshoot). We wondered whether this is due to sensory or motor fatigue, defined as the molecular or physical incapacity to trigger or activate a response (Rankin et al., 2009; Thompson, 2009). In other words, are roots losing the competence for electrotropic response after about 3 hours of exposure? For example, if damaged or depleted of key signaling molecule (other than *tZ*)? Alternatively, is it a case of desensitization, or habituation, where the response to the same continuous stimulus decreases in time without any sensory or motor fatigue (Rankin et al., 2009)? We reasoned that one way to answer this would be to wait until roots settled on a stable ESA and then suddenly change the EF orientation to test whether they are still able to respond to that type of stimulus. We exposed the roots to +1.5 V/cm for 17 h before inverting the EF orientation to the opposite direction with -1.5 V/cm at 17 h. Strikingly, roots responded to this change immediately by turning again towards the negative electrode, now at the opposite side of the V-box because of the EF inversion (Figure 3, a; t-test between 17 h and 20 h, p < 0.001), therefore ruling out the fatigue hypothesis. To confirm that the roots were still viable and able to respond to gravity, we switched the EF off 5 h after the inversion and observed the expected gravitropic response (Figure 3, a).

**Figure 3.**
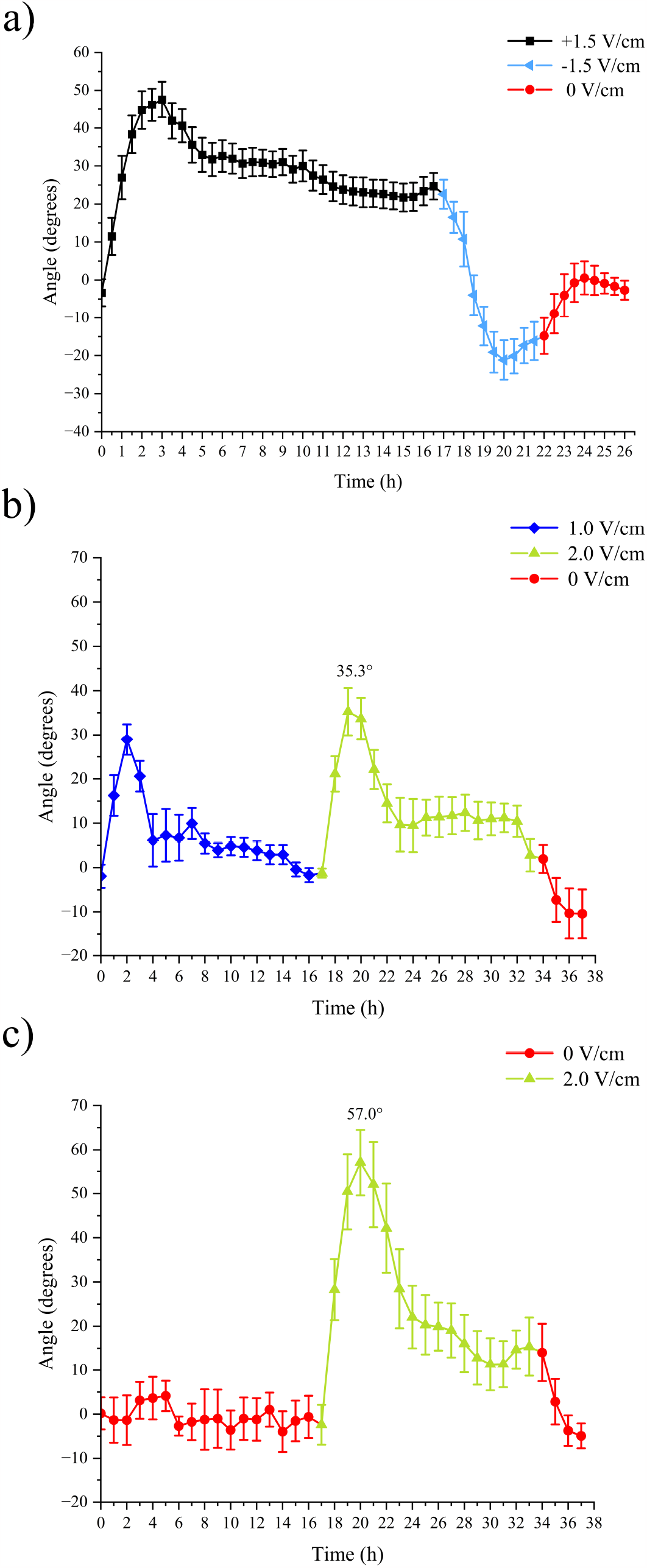
Habituation and Hysteresis. Average WT root tip orientations relative to the gravity vector versus time of plants exposed to sequences of EFs. **a**, +1.5 V/cm → -1.5 V/cm → 0 V/cm (N=16, R=5); **b**, 1.0 V/cm → 2.0 V/cm → 0 V/cm (N=9, R=2); **c**, 0 V/cm → 2.0 V/cm → 0 V/cm (N=10, R=2). See main text for statistical analysis. N, sample size; R, number of replicates. Error bars, s.e.m.

To further confirm that the relaxation and stable ESA were not due to fatigue, we devised an alternative, independent, way to induce a new response after the first ESA was established. We exposed the roots to 1.0 V/cm for 17 h before increasing the EF to 2.0 V/cm for a further 17 h. As in the previous scenario, we found that as soon as the EF was changed, roots responded again with an overshoot, followed by relaxation towards a new ESA (Figure 3, b; t-test between first ESA at 16-17 h and second overshoot at 19 h, p < 0.001). To confirm that the roots were still viable and able to respond to gravity, we switched the EF off after the second ESA and observed the expected gravitropic response (Figure 3, b).

Taken together, these results indicate that the decrease in electrotropic response (relaxation) observed after the first few hours of EF exposure is not a consequence of fatigue but is, instead, an example of habituation (Rankin et al., 2009) and that either an inversion or an increase of EF is sufficient to induce dishabituation.

We then wondered whether a decrease, instead of an increase, of the EF would also be sufficient for dishabituation. We applied 2.0 V/cm for 17 h, followed by a further 17 h at 1.0 V/cm. We did not observe a second response in this case (Supplemental Figure S2; t-test between maximum at 3 h and average root tips angle at 19-21 h, p < 0.001). The fact that only an increase, and not a decrease, in the stimulus intensity is sufficient to trigger a new response is considered a common characteristic of habituation mechanisms (Rankin et al., 2009).

### Hysteresis in long-term electrotropism

As described above, habituated Arabidopsis roots responded to an EF increment from 1.0 to 2.0 V/cm with a second overshoot. Strikingly, although we had previously shown that the short-term root electrotropic response (overshoot maximum) increases as a power of the stimulus intensity (Salvalaio et al., 2022), we noticed that in this experiment the second overshoot at 2.0 V/cm is statistically indistinguishable from the first one at 1.0 V/cm (Figure 3, b; t-test between first and second maxima at 2 h and 19 h, p = 0.3381). To explain this apparent contradiction, we wondered whether previous exposures to the EF had any effect on subsequent responses to the same kind of stimulus, even when at higher intensities.

To address this, we compared the electrotropic response of roots with different histories: those first exposed to 1.0 V/cm, habituated, and then exposed to 2.0 V/cm (Figure 3, b) with those exposed to a 2.0 V/cm EF without any previous experience of EF (Figure 3, c). Remarkably, we found that the overshoot angle at 2.0 V/cm in the two different scenarios is different (t-test between the two maxima at 2.0 V/cm in 3b and 3c, p < 0.05). In other words, pre-exposure to 1.0 V/cm affected the response to 2.0 V/cm, proving that the entity of the electrotropic response does not depend on the absolute intensity of the stimulus but on its step-increase.

Taken together, these results suggest that the entity of the electrotropic response depends on the history of the system, which denotes a process of hysteresis or “stress imprint” (Bruce et al., 2007).

### Spontaneous dishabituation

A defining characteristic of habituation is that it is maintained for a finite amount of time even after the original stimulus is removed (Peretz, 1970; Peters et al., 2007; Rankin et al., 2009). In other words, once the organism has habituated to a stimulus, when this is turned off it will take some time for the organism to be able to respond again to a new instance of the same stimulus (spontaneous dishabituation). The amount of time elapsed between the interruption of the stimulus and the spontaneous dishabituation is an important cue to the molecular mechanism underlying habituation and, in this case, electrotropism.

To characterize this, we first induced habituation by exposing the roots to 1.5 V/cm for 10 h and then paused the EF for different periods before switching it on again at 1.5 V/cm (Figure 4). A reappearance of response (roots turning again towards the negative electrode) would indicate that the habituation had been spontaneously erased. A 0.25 h pause was not sufficient to allow spontaneous dishabituation (Figure 4, a; t-test between 10.25 and 15.5 h, p = 0.101), while a 0.5 h pause already resulted in a partial dishabituation, where a significant response is detectable but the new maximum is not comparable in magnitude to the first one (Figure 4, b; t-test between 10.5 h (first ESA) and 13 h (2^nd^ maximum), p < 0.01; t-test between 1^st^ and 2^nd^ maxima, p < 0.001). We found that a 1 h pause was already sufficient to fully recover the electrotropic response (Figure 4, c; t-test between 1^st^ and 2^nd^ maxima, p = 0.4155), indicating that the minimum pause required to induce a full dishabituation is between 30 min and 1 hour.

**Figure 4.**
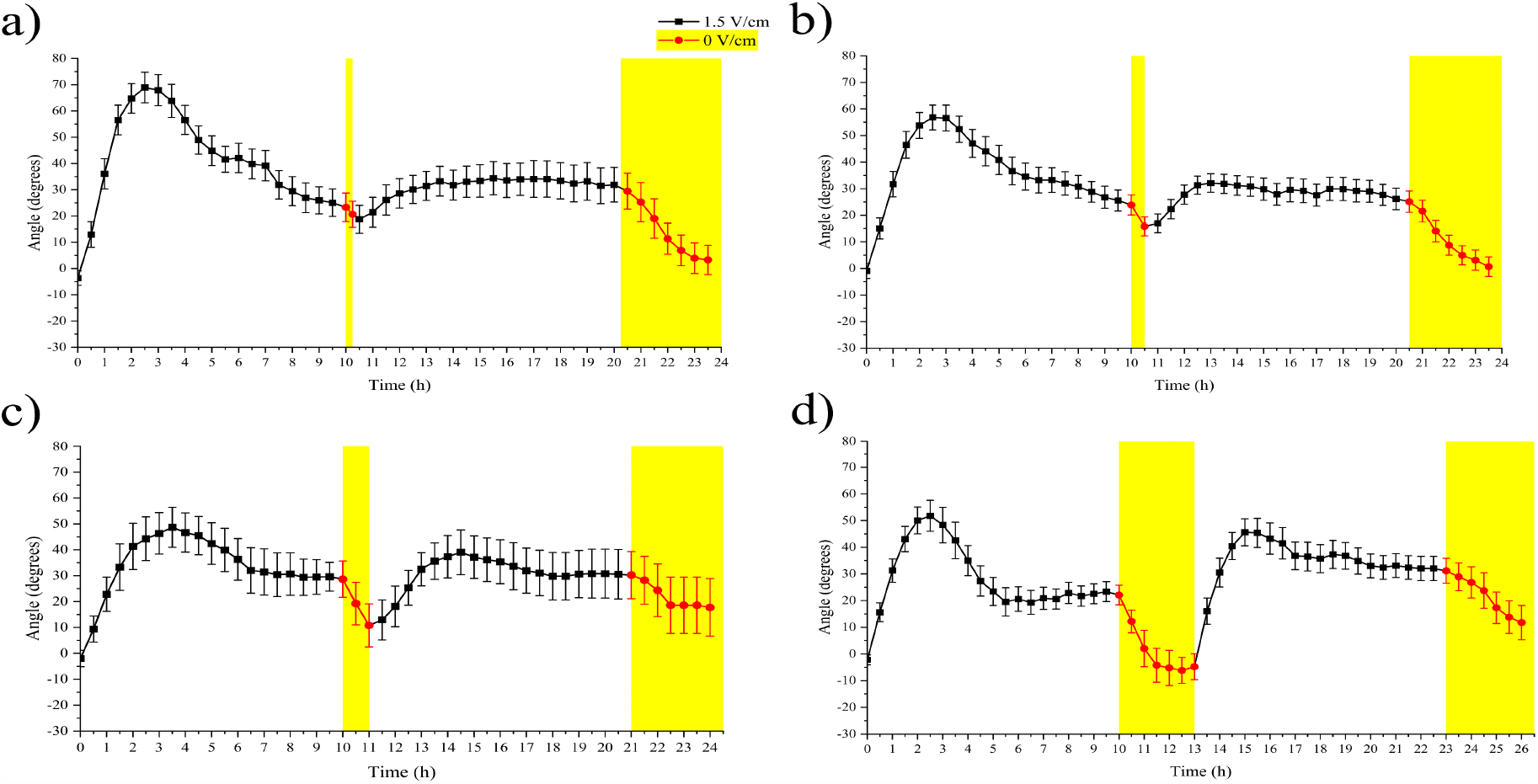
Spontaneous dishabituation. Average WT root tip orientations relative to the gravity vector versus time when a pause of 0 V/cm (red line and yellow background) is applied. **a**, 0.25 h, N=13, R=3; **b**, 0.5 h pause, N=24, R=5; **c**, 1.0 h pause, N=10, R=2; d, 3.0 h pause, N=18, R=4. See main text for statistical analysis. N, sample size; R, number of replicates. Error bars, s.e.m.

We noticed that, as expected, roots responded to gravity as soon as the EF was turned off, but it took them at least 3 hours to fully align vertically (Figure 4, d).

Finally, when we implemented a 3 h pause and allowed the roots to fully reorient vertically, the response to the new 1.5 V/cm was even more dramatic and the overshoot truly resembled what is observed in roots that were never exposed to the EF (Figure 4, d; t-test between 1^st^ and 2^nd^ maxima, p = 0.5010).

## DISCUSSION

This work extends our previous quantitative characterization of root electrotropism (Salvalaio et al., 2022) to describe the long-term behavior of roots exposed to external electric fields. It is a necessary step towards a full understanding of the molecular and physical mechanisms controlling electrotropism in plants.

Here, we have shown that the long-term root electrotropic response is more complex than expected, starting with a quick bending of the root tip towards the cathode of the electrolytic cell (our V-box and V-slide), followed by a slower relaxation phase when the root bends downwards to settle on a steady-state orientation at a non-zero angle with respect to gravity (Figure 1, a). The analogy with common response curves observed in other living systems is evident, and we adopted some of the terms to describe the phenomenon. We call the maximum angle reached in the first 2-3 hours an “overshoot” and the final stable angle an “Electrotropic Set-point Angle”, or ESA.

Since every effort was made to perform these experiments in a controlled environment with constant and uniform temperature, illumination, and chemical composition, the two dominant stimuli were from the electric field (or the consequent ionic current established in the medium) and the gravitational field. Both are perceived by the root at any given time, and the orientation of the root must be interpreted as the combination of these two competing tropisms. We had previously shown that auxin asymmetric distribution is not necessary for root electrotropism (Salvalaio et al., 2022), and that “decapitated” roots missing the entire root cap (with amyloplasts) and the meristematic zone up to 400 µm from the root tip, are still strongly electrotropic although obviously agravitropic (Salvalaio et al., 2022), suggesting that the two tropisms act through independent mechanisms. Moreover, although the characteristic Gravitropic Set-point Angle (GSA) of lateral roots has been correlated with asymmetric distribution of cytokinin (Waidmann et al., 2019), here we have shown that cytokinin, although necessary for electrotropism (Salvalaio et al., 2022), is not asymmetrically distributed during electrotropism (Figure 1, c and d). Taken together, these results suggest that the observed ESA is not just a variation of the GSA and that the two tropisms are truly distinguished.

The fact that the ESA is usually larger than zero, where zero degrees is the orientation parallel to the gravity vector, indicates that electrotropism is not completely abolished by longterm exposure to the stimulus, but only significantly inhibited. Interestingly, we had previously reported that the response curve for early root electrotropism, which now we know is the overshoot response, is a power function of the imposed EF (Salvalaio et al., 2022), while here we show that the final ESA is a linear function of the same EF (Figure 2, a). This suggests that the relaxation phase, or the mechanism leading to habituation (see below), must act as a nonlinear function of the EF that “rectifies” the early response curve into the late one.

One of the most striking results of this work is certainly the discovery of habituation in root electrotropism. The term is borrowed from well-known behavior studies (Thompson and Spencer, 1966; Thompson, 2009) and simply refers to the phenomenon where the progressive decrease of a response results from repeated stimulation but does not involve a loss of competence in sensing or acting on the stimulus (sensory or motor fatigue) (Rankin et al., 2009). Indeed, we have shown that roots that lost the maximum orientation reached during the overshoot and settled to a low ESA, are still perfectly competent to respond to EFs: a sudden inversion of EF direction (Figure 3, a) or increase of its intensity (Figure 3, b) are each sufficient to trigger a new strong electrotropic response. The immediate response of habituated roots when exposed to a new realization (inverted or stronger) of the same type of stimulus (EF) seems to exclude saturation of receptors or other kinds of sensors, depletion of key signaling molecules or simply a loss of cell expansion competence; all examples of sensory and mechanical fatigue in this context. The emergence of habituation in root electrotropism is interesting because it offers a glimpse of the complexity of the largely unknown mechanism behind this tropism.

The fact that a prolonged pause of the stimulus will eventually erase habituation and make the system responsive again is common and describes habituation as a reversible process (Rankin et al., 2009). But what is interesting is that there is a minimum amount of waiting time, without the stimulus, before this dishabituation occurs. In other words, the system generally maintains its “habituated” state for some time, even after the stimulus is turned off. This remarkable characteristic, which we observed in root electrotropism (Figure 4, a), implies a mechanism with some kind of “inertia” or “memory” of the effects of the previous sustained exposure to the stimulus. This is also supported by our results revealing hysteresis in root electrotropism: roots exposed for the first time to 2.0 V/cm respond differently to those previously exposed to 1.0 V/cm, habituated, and then exposed to the same 2.0 V/cm (Figure 3, c vs b). Such dependence on the previous history of the system is also reminiscent of “stress priming” or “memory” (Hilker and Schmülling, 2019) and should be included in any future model describing plant electrotropism.

Finally, although we are still far from being able to draw such a model, a few hypotheses can be made based on the results from this work and others. Ultimately, like any other tropism, some kind of asymmetry between the root’s side facing the negative and the positive electrodes must be established, either at the molecular or physical level. We had shown that cytokinin biosynthesis is necessary for root electrotropism (Salvalaio et al., 2022) but here we show that this hormone does not seem to be asymmetrically distributed during the tropic response (Figure 1, c and d). This indicates that something else must be localized or activated/deactivated in an asymmetric way.

Could other phytohormones be involved in electrotropism? An important candidate is certainly abscisic acid (ABA), which has been shown to play a role in the microtubule reorientation driving halotropism (Yu et al., 2022) and to control root hydrotropism within the cortex layer (Dietrich et al., 2017).

We have also shown some cells in the cortical layer facing the positive electrode expanding dramatically and eventually bursting when exposed to higher EF (Figure 2, c). This suggests that the root cortex might play a key role in the sensing and/or mechanical response to external EFs, making the hypothesis of a role for ABA even more intriguing. Our results also suggest that, while the bending towards the negative electrode is the proper electrotropic response, the counter-bending toward the positive is just a mechanical recoil once enough cells burst on the side facing the positive electrode.

Interestingly, cell wall acidification is thought to be sufficient to trigger cell growth, an idea often referred to as “acid growth theory” (Rayle and Cleland, 1970) and, more recently, this phenomenon has been linked to vacuole expansion (Dünser and Kleine-Vehn, 2015). Although cell wall acidification is usually attributed to auxin through the action of plasma membrane H+-ATPase pumps (Rayle and Cleland, 1992), it is worth considering the hypothesis that an external EF could move protons through the apoplast independently of auxin, resulting in acidification of one side of the tissue (predictably, the one facing the negative electrode). In fact, such an electrophoresis effect doesn’t have to be limited to protons, and could potentially result in asymmetric distribution of other charged molecules on the root surface.

Finally, another, related, possibility is that spatial patterns of plasma membrane potentials could be directly or indirectly perturbed by the external electric field, leading to asymmetric growth. The link between bioelectric fields, cell growth and developmental processes such as regeneration has been widely studied in animals (Levin, 2021) and it is not unlikely to play a role in plant development as well where, for example, external EFs can enhance root regeneration (Kral et al., 2016).

Overall, it is clear that we are just starting to understand the curious phenomenon of electrotropism in plant, and we encourage future work to attempt the isolation of the key genetic and epigenetic regulators of the rich behavior described here.

## MATERIALS AND METHODS

### Plant material and media

*Plant Material*. Wild-type *Arabidopsis thaliana* seeds were from the Columbia (Col-0) ecotype. *TCSn::GFP* seeds were obtained from the Nottingham Arabidopsis Stock Centre (NASC, N69180). In all cases, seeds were imbibed in water and kept in the dark at 4° C for 2 days, to synchronize germination. Seeds were surface sterilized with 50% Haychlor bleach and 0.0005% Triton X-100 for 3 minutes and then washed with sterilized Milli-Q water 6 times.

After sterilization, seeds were sown in PCR tubes containing 1 X MS solid medium. The PCR tubes, whose ends were cut to allow the growing roots to come out, were placed in a 3D-printed holder placed inside Magenta boxes (Sigma-Aldrich V8380) filled with 1/500X MS liquid medium, as previously described (Salvalaio et al., 2022). These Magenta boxes (“nurseries”) were then transferred to a growth chamber at 22°C, with a 16 h/8 h light/dark photoperiod and light intensity 120 µmol m^-2^ s^-1^.

*Media composition*. “1X MS solid”: 0.43% (w/v) Murashige and Skoog (MS) Basal medium (Sigma-Aldrich, M5519), 0.5% (w/v) sucrose (Sigma-Aldrich, S0389), 0.05% (w/v) MES hydrate (Sigma-Aldrich M8250), 0.8% (w/v) agar (Sigma-Aldrich 05040), pH adjusted to 5.7 with Tris Buffer (Fisher-Scientific 10205100). “1/500X MS liquid”: 0.00088% (w/v) Murashige and Skoog (MS) Basal medium (Sigma-Aldrich, M5519), 0.5% (w/v) sucrose (Sigma-Aldrich, S0389), 0.05% (w/v) MES hydrate (Sigma-Aldrich M8250), pH adjusted to 5.7 with Tris Buffer (Fisher-Scientific 10205100).

### Electrotropism assay (V-box)

Seedlings of Col-0 were grown in nurseries as described in the “Plant material” section. PCR tubes containing 7-day-old seedlings were transferred to a 3D-printed holder made to hold up to 5 seedlings and platinum-iridium foil electrodes (Platinum:Iridium = 80:20; Alfa Aesar 41805.FF), as previously described (Salvalaio et al., 2022). The holder containing the seedlings was placed inside a Magenta box, renamed “V-box”, filled with 150 ml of 1/500X MS liquid medium. The rear body of the V-box was perforated with two holes to allow the wires attached to the top of the electrodes, later connected to an external power supply, to exit the box. The lid of the V-box was also customized to create a sealed environment, which was crucial to perform long-term electrotropism experiments in total sterility. Precisely, a 4-port screw cap (Fisher Scientific 15710239, 10784724, 10689163) was attached with silicone paste to the top of the original Magenta box lid, previously drilled in correspondence with the four ports. Silicone tubings were run through the ports inside the V-box and a bottle containing 3600 ml of 1/500X MS liquid medium. The continuous flow of medium through the system was powered by peristaltic pumps (Verdeflex AU R2550030 RS1), as previously described (Salvalaio et al., 2022).

The electric field was generated using a time-programmable power supply (B&K precision BK 9201), which was attached to electric wires soldered to the two electrodes of the V-box, always kept outside the medium to avoid contaminants from the solder. The electrodes were positioned to generate an electric field perpendicular to the gravity vector, i.e. horizontal.

We report a “nominal” electric field (EF, in V/cm), simply defined as the electric potential difference maintained between the two electrodes, divided by their distance. This corresponds to the electric field that would be generated in vacuum. Given the notorious complexity of the double-layer effect at the surface between electrodes and liquid medium, a full quantitative description of the electric field generated inside the medium would be beyond the scope of this paper. Instead, we calculate the current density *D*_*i*_ = *I*/*A*_*e*_ (where *I* is the total current intensity measured in the circuit and *A*_*e*_ is the surface area of each electrode) and report the current impacting the root (or the corresponding cross-section of the current density) *I*_*r*_ = *D*_*i*_ × *A*_*r*_ (where *A*_*r*_ is the estimated area of the root surface facing each electrode).

The power supply was programmed accordingly to generate a constant EF (long-term electrotropism), a sequence of variable (habituation and hysteresis experiments) and intermittent (spontaneous dishabituation) EF intensities.

The roots were imaged with a Raspberry Pi camera module (V2 913-2664) connected to a Raspberry Pi 4 model B computer (RS 182-2096). Photos were taken every 30 minutes, using the command *crontab* in the local Raspbian OS. To image the roots throughout the experiments, all experiments were performed in a growth room with constant light of 100 µmol m-2 s-1), at 22°C. The images were then analyzed with ImageJ, using the line tool to measure the root tip angle relative to gravity (Figure S1).

WT experiments with continuous, constant EFs were performed with the following sample sizes N and number of replicates R: 0 V/cm, N=19, R=4; 1.0 V/cm, N=34, R=7; 1.5 V/cm, N=77, R=18; 2.0 V/cm, N=50, R=11; 2.5 V/cm, N=29, R=6; 3.0 V/cm, N=30, R=6; +1.5 V/cm to -1.5 V/cm, N=16, R=5. WT experiments with a sequence of different EF intensities were performed with the following sample sizes N and number of replicates R: 1.0→2.0→0 V/cm, N=9, R=2; 2.0→1.0→0 V/cm, N=18, R=4; 0→2.0→0 V/cm, N=10, R=2. WT experiments with discontinuous EFs were performed with the following sample sizes N and number of replicates R: 1.5 V/cm with 0.25 h-break, N=13, R=3; 1.5 V/cm with 0.5 h-break, N=24, R=5; 1.5 V/cm with 1 h-break, N=10, R=2; 1.5 V/cm with 3 h-break, N=18, R=4.

### Electrotropism on a microscope (V-slide)

Seedlings of Col-0 and *TCSn::GFP* were grown in nurseries as described in the “Plant material” section. After 7 days, the seedlings were transferred to a 3D-printed modified microscope slide (V-slide), where two platinum-iridium foil electrodes were used to impose EFs on roots to be imaged with the confocal microscope immediately afterwards, as previously detailed (Salvalaio et al., 2022).

An electric field of 3.0 V/cm, which corresponded to an Ir of 0.48 ± 0.10 mA (defined in the V-box section of Materials and Methods), was generated by connecting the two electrodes of the V-slide to a power supply (Tenma 72-10495). Throughout the experiments, 100 ml of 1/500X MS liquid medium was circulated through a system of tubing and peristaltic pumps (Boxer 9K, 9016.930), at a flow rate of 1.66 ml/min. During the last 10 minutes of every experiment, roots were counter-stained with 10 µg/ml Propidium Iodide (PI) (Sigma Aldrich P4170), which was added directly to the medium of the V-slide. Subsequently, after turning off the EF and the circulating system, the staining solution was removed using a pipette, and the roots were covered with a coverslip. Lastly, roots were analyzed under an inverted Leica SP5 confocal laser scanning microscope, with a 20X air objective. GFP (for experiments with *TCSn::GFP*) and PI (for experiments with Col-0 and *TCSn::GFP*) were excited with the 488 nm Argon laser line and the emission collected with PMT detector at 500-570 nm (GFP) and 600-700 nm (PI).

Confocal imaging on V-slide experiments was performed with the following sample sizes N and number of replicates R: Col-0, 0 V/cm (0 mA) N=14, R=3; 3.0 V/cm (0.48 ± 0.10 mA) N=42, R=12. *TCSn::GFP*, 0 V/cm (0 mA) for 10 min, N=14, R=3; 3 V/cm (0.48 ± 0.10 mA) for 10 min, N=17, R=6; 3.0 V/cm (0.48 ± 0.10 mA) for 30 min, N=14, R=5; 3.0 V/cm (0.48 ± 0.10 mA) for 45 min, N=24, R=7.

### Fluorescence intensity measurement

*TCSn::GFP* seedlings were stained with propidium iodide to visualize root anatomy and identify the quiescent center (as described in “Electrotropism on a microscope (V-slide)”). The GFP fluorescence intensity was measured with ImageJ. In detail, a segmented line (width=25) was drawn along the lateral root cap cell file, within 200 µm above the quiescent center as described in similar protocols using the same reporter (Chang et al., 2019). We then measured the average pixel fluorescence intensity on the line. For each root, the fluorescence intensity measured on the side of the root facing the negative electrode was subtracted from the one measured on the side facing the positive electrode, and the number obtained was subsequently divided by the total fluorescence. This “symmetry parameter” is expected to be zero for symmetric distributions. For the control at 0 V/cm, the fluorescence intensity on either the right-or left-hand side of the root was subtracted randomly, to get an unbiased ratio.

### Cytokinin treatment

Col-0 seedlings were grown in nurseries (as described in “Plant material”) where both the 1X MS solid and the 1/500X MS liquid media were supplemented with 10 nM *trans*-zeatin (Sigma-Aldrich, Z0876). After 7 days, PCR tubes containing the seedlings were transferred into V-boxes filled with 1/500X MS liquid + 10nM *t*Z. The electrotropism experiments were run at 1.5 V/cm for 17h, and the reservoir medium circulated throughout the experiments was also supplemented with 10 nM *t*Z.

WT + 10nM *t*Z experiments were performed with the following sample sizes N and number of replicates R: N=16, R=4.

### Statistical analysis

The Shapiro-Wilk test with alpha level = 0.05 was used to test the normality of the samples. For normal distributions, the two-tails Student’s t-test assuming unequal variances was used to determine if two data sets had significant differences. If one of the two distributions was not normal, the Mann-Whitney test was used.

One-way ANOVA at the 0.05 level was used when there were more than two data sets to compare.

In the fluorescence intensity quantification, one sample t-test with alpha level = 0.05 was used to test whether the mean of the population was equal to 0. All statistical tests were performed with Excel and OriginPro.

## ACKNOWLEDGEMENTS AND FUNDING

We thank M. Schwarze for helping with the initial literature survey on habituation. This work was partly supported by Imperial College’s Excellence Fund for Frontier Research.

