## Supplemental Figures 1-2 for "Long-term root electrotropism reveals habituation and hysteresis"

### Supplemental Information

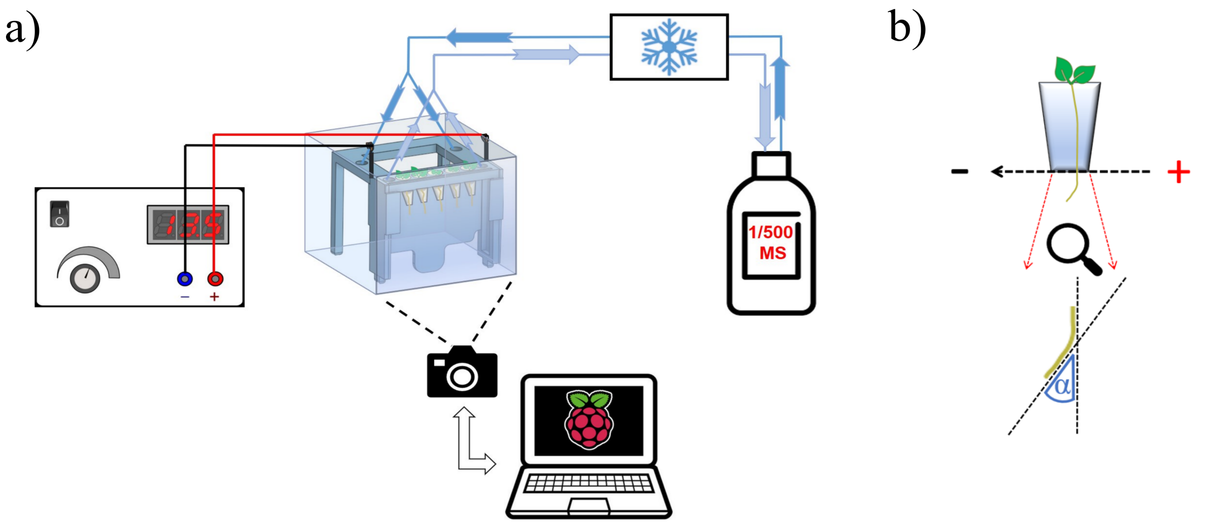


**Supplemental Figure 1.** Root electrotropism assay. **a**, Schematic of the V-box connected to the power supply and to the medium circulation system; **b**, measured angle between the root tip and the gravity vector.

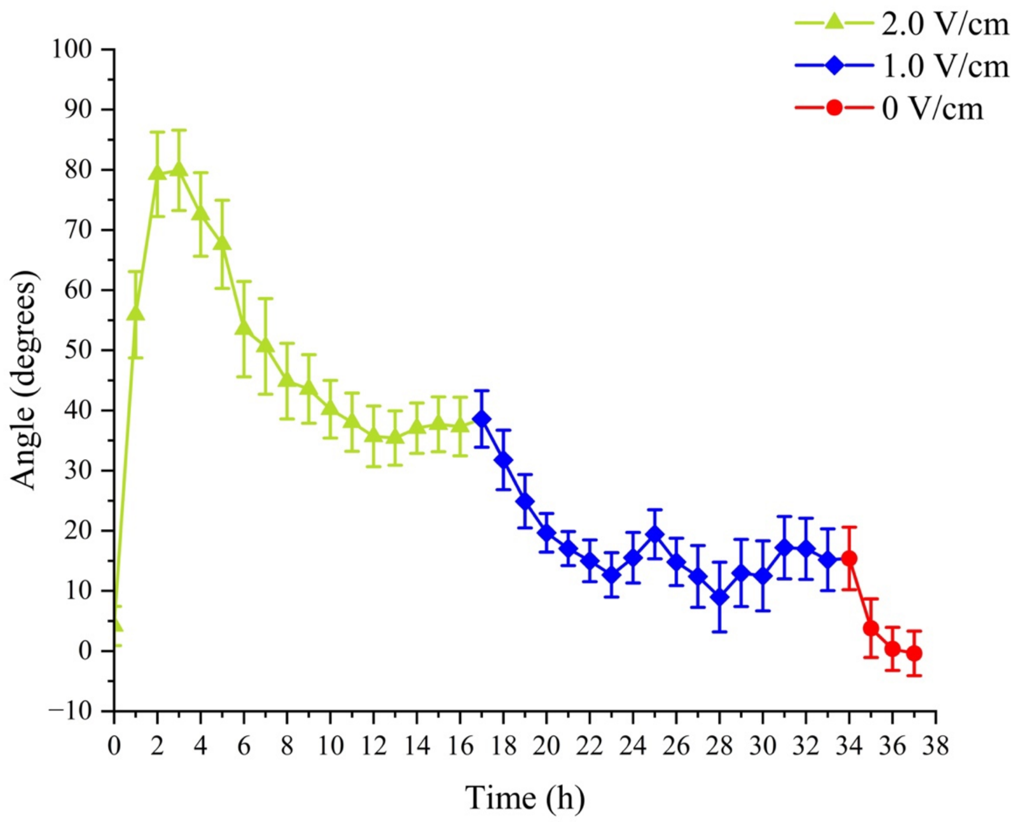


**Supplemental Figure 2.** Decrease of E-field intensity. Average WT root tip orientations relative to the gravity vector versus time when a change in EF intensity 2.0 V/cm → 1.0 V/cm → 0 V/cm (N=10, R=2) is applied; N=18, R=4. See main text for statistical analysis. N, sample size; R, number of replicates. Error bars, s.e.m.
